# Investigating the viral ecology of global bee communities with high-throughput metagenomics

**DOI:** 10.1101/243139

**Authors:** David A. Galbraith, Zachary L. Fuller, Axel Brockmann, Maryann Frazier, Mary W. Gikungu, Karen M. Kapheim, Jeffrey T. Kerby, Sarah D. Kocher, Oleksiy Losyev, Elliud Muli, Harland M. Patch, Joyce M. Sakamoto, Scott Stanley, Anthony D. Vaudo, Christina M. Grozinger

## Abstract

Bee viral ecology is a fascinating emerging area of research: viruses exert a range of effects on their hosts, exacerbate the impacts of other environmental stressors, and, importantly, are readily shared across multiple bee species in a community. However, our understanding of bee viral communities is limited, as it is primarily derived from studies of North American and European *Apis mellifera* populations. Here, we examined viruses in populations of *A. mellifera* and 11 other bee species from 9 countries, across 5 continents and Oceania. We developed a novel pipeline to rapidly, inexpensively, and robustly screen for bee viruses. This pipeline includes purification of encapsulated RNA/DNA viruses, sequence-independent amplification, high throughput sequencing, integrated assembly of contigs, and filtering to identify contigs specifically corresponding to viral sequences. We identified sequences corresponding to (+)ssRNA, (-)ssRNA, dsRNA, and ssDNA viruses. Overall, we found 127 contigs corresponding to novel viruses (i.e. previously not observed in bees), with 29 represented by >0.1 % of the reads in a given sample. These viruses and viral families were distributed across multiple regions and species. This study provides a robust pipeline for metagenomics analysis of viruses, and greatly expands our understanding of the diversity of viruses found in bee communities.

## Introduction

Populations of bees around the world are exhibiting declines, which are the result of multiple interacting factors, including pressure from pathogens [1]. To date, more than 24 viruses have been identified in western honey bees (*Apis mellifera*) [2–7]. These viral infections can result in a range of symptoms, from no obvious phenotype to rapid death and colony loss, depending on the viral species, viral strain, physiological state of the host, and presence of other stressors (reviewed in [8]). However, despite the importance of bees as pollinators of flowering plants in agricultural and natural landscapes [12,13] and the importance of viruses to bee health, our understanding of bee viruses is surprisingly limited: despite the diversity of bee species and their worldwide distribution, the vast majority of the studies examining bee pathogens have focused on western honey bees (*Apis mellifera*) populations in North America and Europe [14].

Bee viral ecology is particularly complex, since many viruses seem to be shared across diverse bee species (see review from Tehel, Brown & Paxton: [14]). Several studies have demonstrated that viruses can spill-over from managed *A. mellifera* or bumble bee (*Bombus spp*) colonies to wild bee populations, with increased viral prevalence in areas with increased density of infected managed colonies [15–19]. Viruses found to be pathogenic in *A. mellifera* (such as deformed wing virus, Israeli acute paralysis virus, acute bee paralysis virus, and Kashmir bee virus) have also been found to be pathogenic in wild bee species [17,20,21]. Targeted studies have identified subsets of the previously identified honey bee viruses (including deformed wing virus, black queen cell virus, sacbrood virus, Israeli acute paralysis virus, acute bee paralysis virus, Lake Sinai virus, chronic bee paralysis virus, and Kashmir bee virus) in bee populations in Asia [22], Africa [23], the Middle East [24], Australia [25], and South America [26,27], suggesting a world-wide distribution of these viruses.

Since the most well-studied viruses were originally identified in *A. mellifera*, it has often been assumed that *A. mellifera* are the primary hosts and other bees are only secondarily infected. However, because many viruses seem to be so readily shared, it is not possible to determine which bee species, if any, serves as the primary host. Indeed, based on sequence analyses, the viral species that have been examined thus far seem to be shared within a geographic region rather than within a particular bee species, suggesting that there may not necessarily be bee species-specific viral strains [16,17]. Thus, it is likely that non-Apis populations also harbor viruses which can spill over into *A. mellifera*.

Similarly, recent metagenomics studies of bees have identified many viruses that were originally described in plants [28,29]. It has been assumed that these are simply contaminants from the pollen and nectar in the bees’ diets, but these viruses may in fact be using bees as hosts. Indeed, tobacco ringspot virus (TRSV) was recently shown to infect and replicate in *A. mellifera* [30].

Thus, fully understanding the viral community in managed and wild bee species, and the plants they feed on, requires a holistic approach that examines these communities in diverse geographic regions. Distinct types and species of viruses undoubtedly remain unidentified in populations of *A. mellifera* and other bee species across the world. Moreover, due to increased globalization, it is likely only a matter of time before these viruses (and other pathogens and parasites) are spread to new regions. For example, recently a colony of an exotic stingless bee species (*Plebeia frontalis*) was found to be established in California [31], potentially harboring pathogens that could be spread to other bee species within the region.

With the development and optimization of inexpensive high-throughput sequencing technologies, it is now possible to sequence the genetic material of the entire community of viruses present within a host, without any prior knowledge of the viral genome sequences (viral metagenomics analysis) [32]. This technology allows rapid and efficient identification of previously uncharacterized types and strains of viruses, bypassing labor-intensive steps of purifying and culturing the viruses. Viral metagenomics has been utilized by many studies seeking to identify novel viruses and characterize viral communities in diverse organisms (reviewed in [33]).

Using this approach, several recent studies have identified novel viruses in populations of bees (*A. mellifera, Bombus terrestris, Bombus pascuorum, Osmia cornuta*, and *Andrena vaga*) collected from sites in the Netherlands, South Africa, Tonga, Israel and Belgium [6,29,34]. However, while these studies have provided valuable information about viral communities within particular species or regions, they are limited in scope, in that they focus on a single bee species in multiple locations (*A. mellifera* populations in Israel [34] or the Netherlands, South Africa, and Tonga [6]) or a selection of wild bees in a single region (Belgium [29]). Furthermore, two of these studies [6,29] did not purify viral particles prior to sequencing, and thus had samples with a high percentage of reads corresponding to the host bee species, which can limit detection of rare viral species.

To improve our ability to detect viruses, we have developed a protocol for unbiased, sequence-independent detection of novel viruses within diverse bee populations from around the globe. We used this approach to characterize the viral communities of 12 bee species collected from 9 countries across 5 different continents and Oceania (see Table 1 for bee species and their respective locations). This approach allowed us to search for novel viruses, begin to determine the distribution of these viruses (and viral families) across the world, and begin to evaluate the extent to which *A. mellifera* and wild bee populations share these viruses. As in other studies [28,29], we identified several viruses that share sequence homology to plant viruses and thus may be the result of contamination, but given that these were found in multiple populations and locations, it raises the question of whether or not these viruses may use bees as hosts.

**Table 1.** Sample information

| Sample ID | Location | Species | Original ID | GPS Coordinates (°Lat., °Long.) |
| --- | --- | --- | --- | --- |
| Positive Control | Pennsylvania, USA | *Apis mellifera* | DWV | 40.8198, -77.8588 |
| PA1-Am | Pennsylvania, USA | *Apis mellifera* | C21-S2 | 40.8369, -77.8786 |
| PA2-Am | Pennsylvania, USA | *Apis mellifera* | C22-S3 | 40.7106, -77.9645 |
| PA3-Am | Pennsylvania, USA | *Apis mellifera* | C23-S4 | 40.7534, -77.6699 |
| PA4-Am | Pennsylvania, USA | *Apis mellifera* | SCAm | 40.8198, -77.8588 |
| PA-Bi | Pennsylvania, USA | *Bombus impatiens* | SCBi | 40.8198, -77.8588 |
| WA-Mr | Washington, USA | *Megachile rotundata* | WSmR | 46.0177, -118.6975 |
| WA-Nm | Washington, USA | *Nomia melanderi* | WSNm | 46.0177, -118.6975 |
| WA-Am | Washington, USA | *Apis mellifera* | WSAm | 46.0177, -118.6975 |
| CA-Am | California, USA | *Apis mellifera* | C24-C1 | 41.7928, -124.1208 |
| NI-Am | Nicaragua | *Apis mellifera* | NIAm | 12.1184, -86.1224 |
| NI-Ts | Nicaragua | *Trigona silvestriana* | NIST | 12.1184, -86.1224 |
| PN-Am | Panama | *Apis mellifera* | PNAm | 9.1500, -79.8500 |
| PN-Mc | Panama | *Megalopta centralis* | PNMg | 9.1500, -79.8500 |
| GE-Am | Georgia | *Apis mellifera* | GAAm | 42.0681, 42.4403 |
| CH-Am | Switzerland | *Apis mellifera* | SWAm | 46.5197, 6.6323 |
| CH-Bs | Switzerland | *Bombus* spp. | SWNb | 46.5197, 6.6323 |
| UA-Am | Ukraine | *Apis mellifera* | UKAm | 50.4501, 30.5234 |
| KE1-Am | Kakamega, Kenya | *Apis mellifera* | C11-G6 | 0.2527, 34.9232 |
| KE2-Am | Kakamega, Kenya | *Apis mellifera* | C6-G1 | 0.2277, 34.8459 |
| KE3-Am | Kisii, Kenya | *Apis mellifera* | C4-K1 | -0.7423, 34.8045 |
| KE4-Am | Kisii, Kenya | *Apis mellifera* | C5-K2 | -0.7399, 34.8895 |
| KE5-Am | Mbita, Kenya | *Apis mellifera* | C16-M2 | -0.4803, 34.2929 |
| KE6-Am | Mbita, Kenya | *Apis mellifera* | KNAm | -0.4354, 34.2144 |
| KE7-Am | Naivasha, Kenya | *Apis mellifera* | C1-N1 | -0.07547, 36.4384 |
| KE8-Am | Naivasha, Kenya | *Apis mellifera* | C2-N2 | -0.5713, 36.3603 |
| KE9-Am | Naivasha, Kenya | *Apis mellifera* | C3-N3 | -0.6616, 36.3835 |
| KE-XY | Mbita, Kenya | *Xylocopa* spp. | KNXY | -0.4734, 34.1823 |
| IN-Ac | UAS-GKVK, Bangalore, India | *Apis cerana* | INAc | 13.0774, 77.5778 |
| IN-Ad | UAS-GKVK, Bangalore, India | *Apis dorsata* | INAd | 13.0774, 77.5779 |
| IN-Af | UAS-GKVK, Bangalore, India | *Apis florea* | INAf | 13.0711, 77.5802 |
| IN-Am | UAS-GKVK, Bangalore, India | *Apis mellifera* | INAm | 13.0712, 77.5805 |
| IN-Ti | UAS-GKVK, Bangalore, India | *Tetragonula iridipennis* | INST | 13.0752, 77.5800 |
| NZ1-Am | Hamilton, New Zealand | *Apis mellifera* | NZN1 | -37.7870, 175.2793 |
| NZ2-Am | Tauranga, New Zealand | *Apis mellifera* | NZN2 | -37.6878, 176.1651 |
| NZ3-Am | Christchurch, New Zealand | *Apis mellifera* | NZS1 | -43.5321, 172.6362 |
| NZ4-Am | Otago, New Zealand | *Apis mellifera* | NZS2 | -44.8280, 169.6345 |

## Results

### Identification of Viral Sequences

We identified 531 different contigs among the 37 samples that shared sequence homology with previously identified viral sequences (see Supplementary Table 2 for information on each of the contigs identified in each sample). Of these, 308 contigs correspond to “novel” viruses (defined as viruses that were not previously found in bees), and 223 contigs correspond to known bee viruses. Many of these contigs were identified in multiple locations and/or species. When duplicate/overlapping contigs are removed, the number of unique novel virus contigs was reduced to 127. Next, we examined the number of contigs that corresponded >0.1% of the total reads for the given sample in which it was found. Only 29 contigs matched these criteria (see Supplementary Table 3).

Sequence homology analyses of these 29 novel virus contigs revealed 9 different viral families among these bee associated viruses, including 6 viral families that had not been heretofore observed in bees (noted in Supplementary Table 3). Of these, one virus from the family *Rhaboviridae* (BRV) was recently described in *A. mellifera* populations in the United States, Israel, South Africa, Tonga and the Netherlands, and *B. impatiens* populations in the United States [6,55]. Of the 29 novel virus contigs, 6 were found to contain a putative RdRp that could be used for phylogenetic analyses.

The distribution and relative quantities of the known viruses and newly identified viruses with RdRp domains (that were present at >0.1% of the reads in each sample) across each sample are shown in Figure 1A. Figure 1B provides information on the geographic distribution of these viruses and their presence in *A. mellifera* samples versus non-*A. mellifera* samples at each site. Details about each of these novel viruses, including their phylogenetic characterization, are presented below. Interestingly, 5 novel viruses were identified in North American samples, 4 in New Zealand, and 2 in each of the other continents; thus, though the North American samples are relatively well-studied compared to the other regions, at least in our analysis, these samples still generated more novel viruses. In regions where both *A. mellifera* and non-*A. mellifera* bees were sampled, *A. mellifera* had higher numbers of viruses in four regions (Kenya, Nicaragua, Switzerland, and Pennsylvania, US), lower numbers in two regions (India and Washington, US) and equivalent numbers in Panama. Only in the United States and India were novel viruses found in non-*A. mellifera* samples that were not present in the *A. mellifera* samples as well. These results suggest that these viruses are globally distributed, both across bee species and geographic regions.

**Figure 1.**
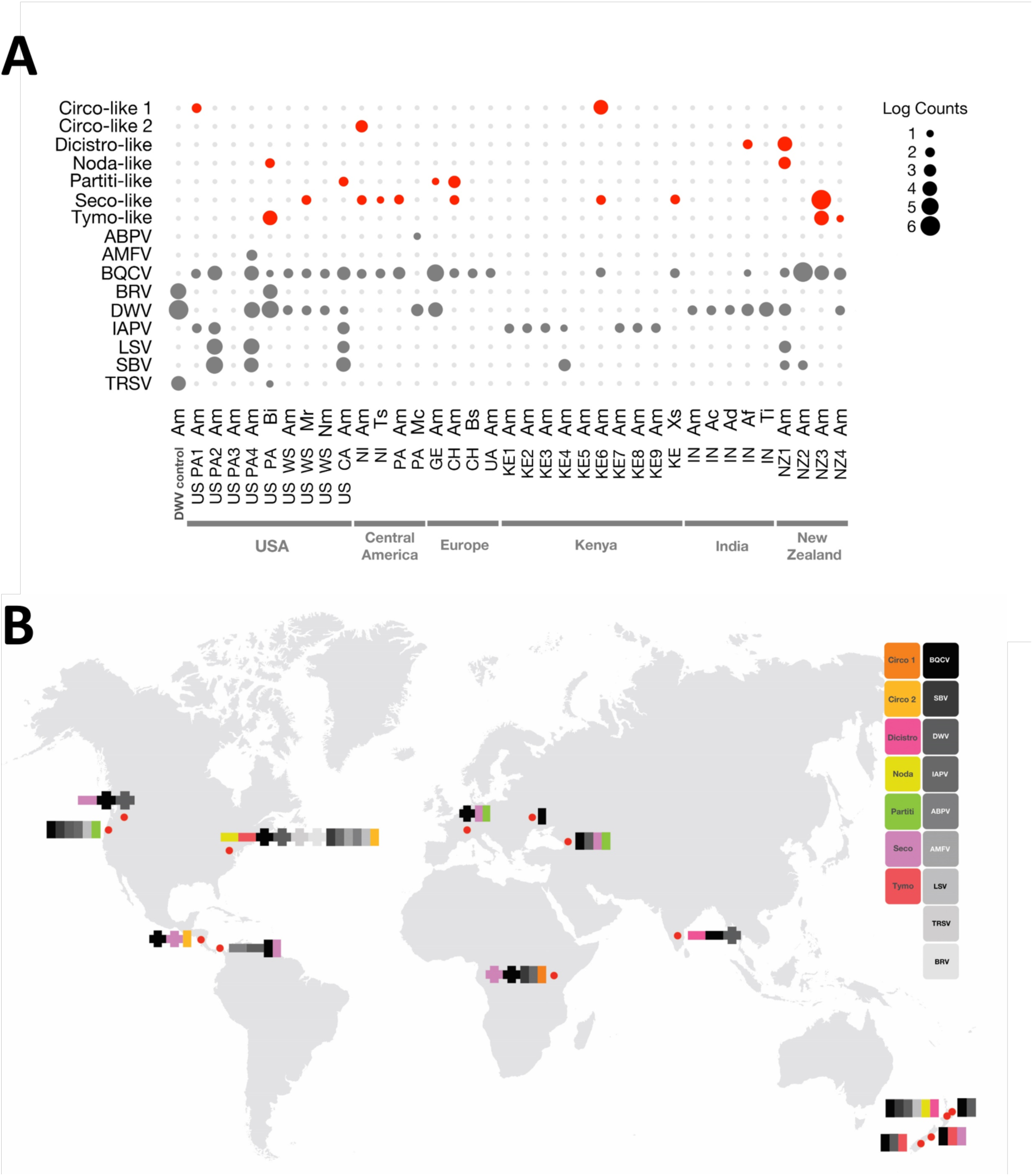
Summary of known and newly identified viral contigs from all samples. These figures summarize the distributions of the known viruses and newly identified viruses (with RdRp sequences and present at >0.1% of the reads in a given sample) from this study. (A) Plot showing the prevalence of all known honey bee viruses (in grey), as well as the 7 novel viruses detected in this dataset that contain an RdRp or other replicase protein (in red) in log read counts. Individual samples are listed according to location, with each sample name containing the location (two letter code for country or, for US samples, state) followed by a two letter code for the species. See Table 1 for complete information on location and species for each sample. (B) This map shows the distribution of known viruses (colored in greyscale) and newly identified viruses (colored) in *A. mellifera* samples (vertical bars) and non-*A. mellifera* samples (horizontal bars) collected at each site. Where a virus was identified in both sample types, the horizontal and vertical bars are overlapping.

Below, we discuss the results for the known viruses, as well as the 6 viruses that possess a putative RdRp.

### Known Viruses

Within this dataset, we identified 7 well-studied bee viruses: black queen cell virus (BQCV), deformed wing virus (DWV), Israeli acute paralysis virus (IAPV), Lake Sinai virus (LSV), acute bee paralysis virus (ABPV), sacbrood virus (SBV), and Apis mellifera filamentous virus (AMFV). BQCV represented the most common known virus among all of the samples, with 22 of the 36 samples testing positive (61%). DWV, another common virus, was detected in 16 (44%) of the samples, and was the second most prevalent virus detected in this study. IAPV (10 samples, 28%), SBV and LSV (6 samples, 17%), and ABPV and AMFV (1 sample, 3%) conclude the known bee viruses identified in our samples (see Figure 1 and Supplementary Table 2 for more details on the locations and species in which these viruses were found).

### Newly Identified Positive Sense RNA Viruses

#### Dicistro-like virus

We identified one Dicistro-like contig that shared sequence homology with known viruses from the family *Dicistroviridae*, a viral family that includes several known bee-infecting viruses, such as IAPV, BQCV, and Aphid lethal paralysis virus (ALPV). This virus was found in *Apis florea* samples from India, with a contig length of 3,818 nt, which included a putative RdRp region, as well as in *A. mellifera* samples from New Zealand with a length of 1,382 nt. A phylogenetic analysis of this region revealed that this particular contig was most closely related to aphid lethal paralysis virus (ALPV), and forms a clade with ALPV and other dicistroviruses identified in a previous study using other invertebrates, including odonates, heteropterans, myriapods, and orthopterans (Figure 2A) [56].

**Figure 2.**
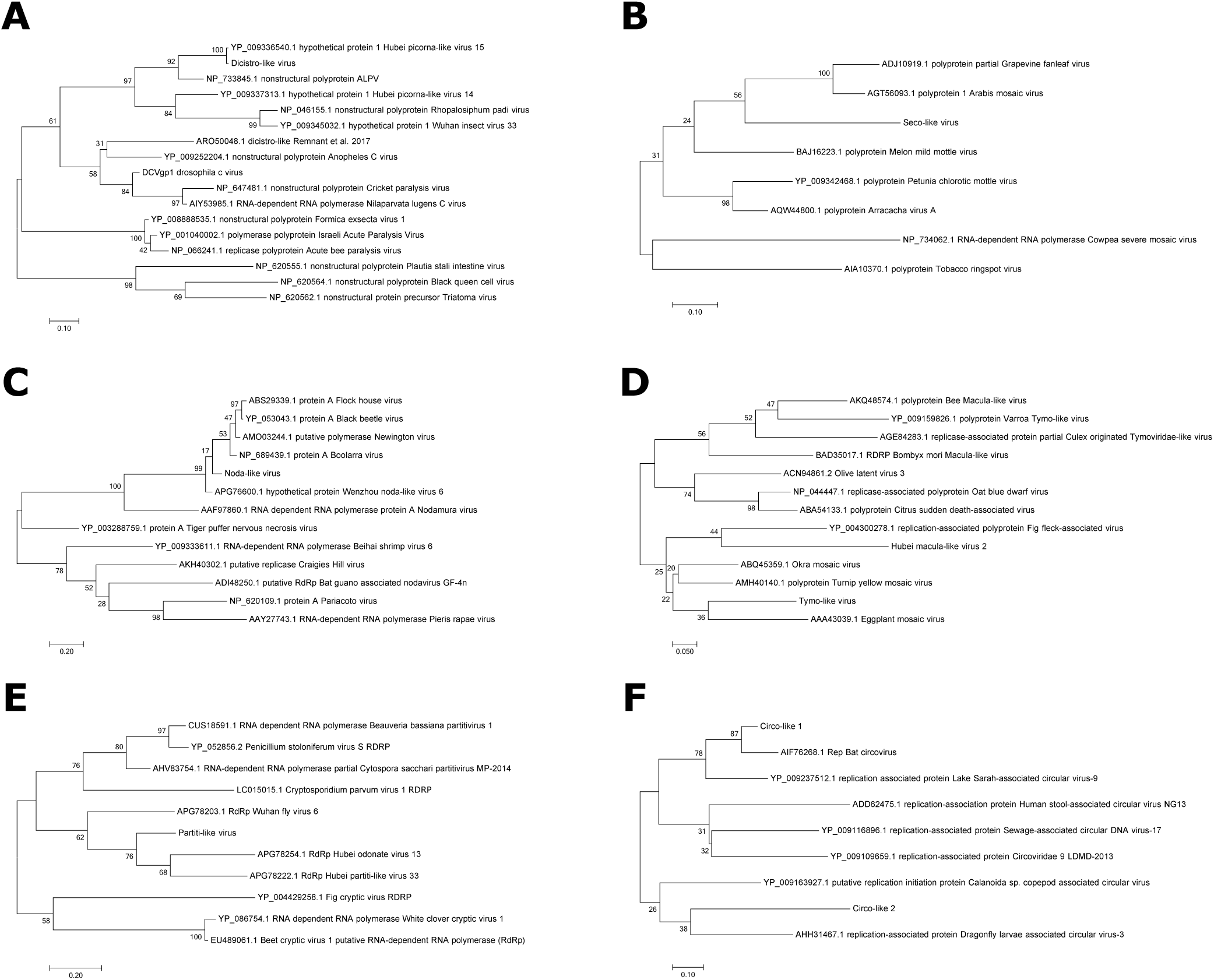
Phylogenetic analyses of novel virus contigs. Maximum-likelihood phylogenetic trees showing the evolutionary relationships between the novel virus contigs containing an RdRp or other replicase protein and other representative sequences from A) *Dicistroviridae*, B) *Secoviridae*, C) *Nodaviridae*, D) *Tymoviridae*, E) *Partitiviridae*, and F) *Circoviridae*. NCBI Accession numbers for the sequences used in these trees can be found in the figure above.

#### Seco-like virus

We identified a Seco-like virus that shares sequence homology to *Secoviridae*, which is in the same viral order as *Dicistroviridae* (Picornavirales). Overlapping viral contigs were found in four bee species across four continents and Oceania (*A. mellifera* in Georgia: 1,988 nt, Kenya: 971 nt, Panama: 1,295 nt, Nicaragua: 1,955 nt, Switzerland: 1,105 nt, and New Zealand: 1,799, *Megachile rotundata* in the US (Washington): 1,500 nt, *Trigona silvestriana* in Nicaragua: 950 nt and *Xylocopa spp*. from Kenya: 1,254 nt). There were minimal differences across the sequences: the putative RdRp region had >99% similarity across the samples. A phylogenetic analysis of the RdRp from this virus suggests close sequence homology to a group of plant viruses (including Arabis mosaic virus; see Figure 2B).

#### Noda-like virus

We identified a novel (+)ssRNA virus within the family *Nodaviridae* in two different species, *A. mellifera* (1,216 nt) and *B. impatiens* (1,831 nt). Interestingly, these were from two distant geographic locations (Hamilton, New Zealand and Pennsylvania, United States). Using a BLASTx [46] approach on the open reading frames identified within the assembled contigs, we identified a putative RdRp that was used for phylogenetic analysis. The phylogenetic analysis revealed that this virus was most closely related to other viruses within the Nodaviridae family of viruses, forming a clade with members of the viral genus Alphanodavirus (Figure 2C), which includes several insect viruses, including flock house virus, black beetle virus, and nodamura virus.

#### Tymo-like virus

A Tymo-like virus with overlapping contigs was identified in *A. mellifera* from two different locations in New Zealand (Christchurch: 6,130 nt and Otago: 276 nt) and *B. impatiens* from Pennsylvania, United States (1,670 nt), sharing sequence homology with the viral family *Tymoviridae*. This viral family is comprised predominantly of plant viruses, however, a recent study also identified a virus from this family in *A. mellifera* samples [5]. The novel virus described here exhibits similar genomic characteristics to other viruses in the Tymovirus genus, with a long poly protein that encodes a methytransferase (MTR), proteinase (PRO), helicase (HEL), and an RNA dependent RNA polymerase (RdRP), a separate overlapping protein that encodes a movement protein, and a coat protein at the 3’ end (see Figure 2D and 3).

**Figure 3.**
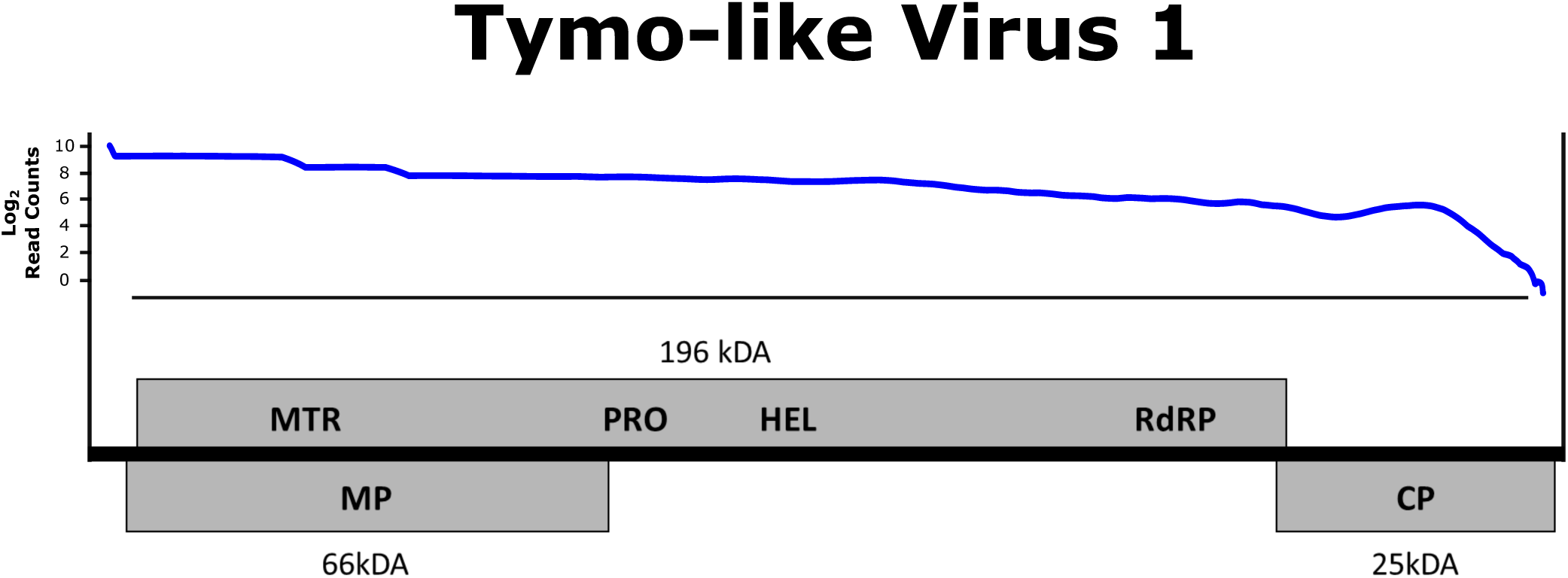
Genome of the newly discovered Tymo-like virus. Genome structure of the Tymo-like virus discovered in *Bombus impatiens* and *Apis mellifera* from the United States and New Zealand. This viral contig contains a long polyprotein encoding a methyltransferase (MTR), proteinase (PRO), helicase (HEL), and an RNA dependent RNA polymerase (RdRP) that is common among viruses from the Tymovirus genus. Additionally, a separate overlapping protein encoding a movement protein at the 5’ end and coat proteins was detected at the 3’ end of the viral contig. Log read counts were calculated using a sliding window of 100 bases.

### Newly Identified Negative Sense RNA Viruses

Only recently have negative sense RNA ((-)ssRNA) viruses been identified in bees [6,29]. One of these, Apis mellifera Rhabdovirus 1 (ARV-1), from the family Rhabdoviridae [6], was detected in *A. mellifera* populations in South Africa, Tonga, and the Netherlands. It was also recently identified in bee populations in Israel and the United States [55]. Our data sets related to this virus were published elsewhere [55] and thus we will only briefly summarize this information here.

We identified contigs corresponding to this Rhabdovirus in two different species (*A. mellifera* and *Bombus impatiens*) that were collected in Pennsylvania, United States, suggesting this virus can easily switch hosts. Thus, Levin et al [55] proposed a different nomenclature, Bee Rhabdovirus (BRV). The two assembled contigs for each species were 13,844 nt and 14,589 nt in *A. mellifera* and *B. impatiens* respectively, and were >99% similar to ARV-1. These assembled contigs contained the 5 genes that are typically carried by Rhabdoviruses, including the large protein (L), a glycoprotein (G), nucleoprotein (N), phosphoprotein (P), and matrix protein (M). Further information about this virus can be found in Levin et al [55].

### Newly Identified Double Stranded RNA Viruses

#### Partiti-like virus

Overlapping contigs with sequence homology to other viruses within the viral family *Partitiviridae* were identified in *A. mellifera* samples obtained from California, United States (1,139 nt), Switzerland (409 nt), and Georgia (385 nt). A phylogenetic analysis of a putative RdRp region from the assembled contigs, determined by a BLASTx [46] screen, revealed sequence homology that is similar to other identified partitiviruses, but most closely related to the newly identified arthropod partitiviruses [56] (Figure 2E).

### Newly Identified Single Stranded DNA viruses

#### Circo-like virus

Sequences corresponding to a circo-like virus in *A. mellifera* from three different locations (Pennsylvania, United States: 521 nt, Kenya: 688 nt, and Nicaragua: 1,531 nt) with sequence homology to viruses from the *Circoviridae* family. Interestingly, a phylogenetic analysis of the replicase protein revealed two different viruses, with the sequences from the US and Nicaraguan *A. mellifera* populations being closely related to circular ssDNA viruses found in bats and dragonflies, respectively (Figure 2F). The Circo-like virus contig that was discovered in Kenya shared sequence homology with bat circovirus (similar to the virus in the Pennsylvania samples), but did not overlap with the replicase protein.

## Discussion

In this study we developed a stringent and robust viral metagenomics pipeline to comprehensively examine the viromes of specimens of 12 different bee species, collected from a broad array of locations (9 countries, across 5 continents and Oceania). This dataset dramatically enhances our current knowledge of bee associated viruses, by identifying 29 novel (heretofore unidentified in bees) viral contigs from 9 viral families that were present at >0.1% of the reads within a sample. Six of these contigs contained a putative RdRp region (identified by BLASTx [46] and ORFfinder[51]) that could be used for phylogenetic analyses to determine relationships between previously identified viruses (see Figure 2). Moreover, using our pipeline, we identified a wide variety of new types of viruses that had not been previously described (or only recently described) in bees, including (-)ssRNA, dsRNA, and ssDNA viruses.

This study represents the largest effort to identify novel pathogens in global bee samples to date. However, it is important to note that this study still had relatively small samples sizes, given the diversity of bee species and their global distributions. Sampling 10 bees per population (as in our study) allows for a 65% probability of detecting viruses with 10% prevalence, and 10% probability of detecting viruses with 1% prevalence [57]. Thus, we have limited ability to identify low-prevalence viruses. Additionally, by focusing on viruses that were represented in >0.1% of the reads in a sample, we may have removed data from low-expressing viruses. Thus, additional screens using larger sample sizes, locations, and species would undoubtedly identify additional viruses and likely demonstrate a broader geographic and species range for the viruses we identified in this study.

It is important to note that we did not assess replication of these viruses in our samples, and this will be necessary to confirm that bees are indeed hosts of these viruses. Viruses may be present in bee samples due to contamination from the environment or their diet; indeed, viruses can infect or contaminate pollen grains and be transmitted to individuals foraging on or consuming this pollen [15,58]. We selected viruses that were present at >0.1% of the reads within a sample (an average of 1,000 reads per sample), to reduce the likelihood that we were examining a trace contaminant. Additionally, all of the viruses were identified in multiple species and multiple sites in independent collections and extractions, and thus it seems somewhat unlikely that these viruses were simply contaminants.

Positive sense RNA ((+)ssRNA) viruses are the most common type of virus that have been identified in bees thus far, with a vast majority of the viruses possessing this genome architecture. Most of these viruses belong to the viral order Picornavirales (including DWV, BQCV, IAPV, and SBV), but recently additionally studies have added to the list of viral orders that infect bees, including Tymovirales [6,29]. Here we have identified novel virus sequences that share sequence homology with three (+)ssRNA virus families, *Dicistroviridae, Secoviridae*, and *Tymoviridae*.

Dicistroviruses are commonly found in honey bee colonies, with the most common viruses being IAPV and BQCV [59]. Unsurprisingly, we found BQCV to be the most widely distributed virus in our dataset, while IAPV was also fairly common, following only behind BQCV and DWV. Another recent study also identified a novel dicistrovirus in *A. mellifera* from Amsterdam, however this virus is phylogenetically different than the virus we described here. We found this viral contig (containing a putative RdRp) in A. *florea* from India, however, we also detected another viral contig from *A. mellifera* from New Zealand that shared sequence homology with a different region of the same virus (Hubei picorna-like virus 15; accession number: NC_032757.1). Unfortunately, there was no overlap between these two contigs, so we cannot say for certain whether these two contigs are a part of the same virus or if they represent two different Dicistroviruses.

We identified a contig corresponding to the Secovirus family in four bee species collected from four continents and Oceania. This virus shares its closest homology with viruses detected primarily in plant species [60]. While additional studies are needed to determine if this virus can replicate in bees, its broad distribution suggests that it is closely associated with bee species. Other recent studies have identified viruses with sequence similarity to plant viruses in bee samples [29,30]. In one case, for tobacco ringspot virus, replication was confirmed in bee samples [30]. It remains to be determined if these viruses are truly plant-associated viruses that simply contaminate bees or if in fact bees are an important host. Regardless, the fact that bees carry plant-associated viruses so frequently suggests that they may be important vectors of plant diseases in the field. Indeed, recent studies found that *A. mellifera* are able to distribute plant viruses through pollination services [61].

A study by de Miranda et al. characterized the first virus from the viral family *Tymoviridae* in *A. mellifera* honey bees, and also found a virus from this family replicating in *Varroa destructor* mites (Bee macula-like virus and *Varroa* Tymo-like virus, respectively) [5]. Our phylogenetic analysis suggested that the virus we observed is distinct from the previously described bee virus in this family, and is more closely related to viruses within the *Tymovirus* genus (as opposed to the *Maculavirus* genus for the other previously identified bee virus in this family). We found evidence of this Tymo-like virus in two distant geographic locations (New Zealand and United States), suggesting that this virus is widespread. Additionally, we found this virus in two different bee species (*Apis mellifera* and *Bombus impatiens*) indicating that this virus can infect both *Apis* and non-Apis bees.

Until recently, none of the viruses found to infect bees were (-)ssRNA viruses. However, Remnant et al. found a novel *Rhabdovirus* in *A. mellifera* honey bees from South Africa, Tonga, and the Netherlands [6]. This group proposed the name Apis mellifera Rhabdovirus (ARV). We also detected this same *Rhabdovirus* (>99% similarity) in *Apis mellifera* and *Bombus impatiens* samples from the US (Pennsylvania). Data on these sequences and detection of the virus in Israel honey bee populations has recently been published [55]. Given that the virus is found in two bee species, we recommend referring to it as Bee Rhabdovirus (BRV).

Partitiviruses have a dsRNA genome, and have been observed mainly in plants and fungi [62]. A recent study examining the invertebrate RNA virosphere [56] identified a number of Partitiviruses within animal hosts from a wide range of arthropod species, including insects. A phylogenetic analysis of our novel *Partitivirus* revealed that it is most closely related to this newly identified group of Arthropod Partitiviruses. The distribution of this virus, as detected here, suggests that it is also widespread, being found in Europe (Switzerland and Georgia) and the US (California). To the best of our knowledge, this is the first *Partitivirus* identified in bees, and further, this is the first evidence of a dsRNA virus in bees.

Viruses that utilize a DNA genome are very rare in bees, with very few being observed to this point [29,63]. Here, we identified a viral contig that is closely related to other members of the *Circoviridae* family of viruses, a viral family that utilizes a circular single stranded DNA (ssDNA) genome. These viruses have been found primarily within birds and mammals, but recent studies have begun to identify circular viruses within insects as well [64], including 24 diverse circular viruses found to be infecting odonates (dragonflies and damselflies) [65]. It is possible that bees harbor a similarly diverse and robust set of circular viruses. The identified Circovirus was phylogenetically related to Dragonfly circularisvirus and to the best of our knowledge, represents the first evidence of a circular ssDNA virus observed in bees.

Our study suggests that, despite the focus on North American and European *A. mellifera* viral populations, these populations still remain to be fully characterized: indeed, the greatest number of newly identified viruses were found in North American samples. Additionally, when considering all the sample sets, it appears that all the newly identified viruses were found in multiple bee species (note that non-overlapping contigs for a novel dicistrovirus found in non-*A. mellifera* samples in India and *A. mellifera* samples in New Zealand – whether these represent distinct viruses or the same virus requires further sequence information). Finally, the viruses were largely evenly shared across the *A. mellifera* and non-*A. mellifera* bee species in a region, though overall, more sites (4 out of 7) had higher numbers of viruses identified in the *A. mellifera* samples than in the non-*A. mellifera* samples.

With our increasing awareness of the importance of viruses to bee health, and the complexity of bee viral ecology – where viruses are shared across insect and potentially plant species – there has been an extensive effort to fully characterize the bee viral community. Prior to 2010, more than 24 viruses were identified in bee populations [2–7]. Since then several additional viruses have been identified using metagenomics approaches [6,29,34]. Together with our results, more than 30 viruses have been found in bees, however, this number likely represents a small subset of the viral diversity present in bee communities. Viral metagenomics for virus discovery in bees has thus far been fruitful, and our protocol has provided a foundation for future studies to continue identifying novel pathogens infecting global bee populations using an inexpensive, unbiased method for the sequence-independent detection of novel viruses. Additional studies are needed to confirm that these viruses replicate within bees, obtain the full viral genome sequences, examine any negative (or positive) impacts infection may have on bees, and examine how these viruses move within bee and plant communities. Our results suggest, however, that viral communities are generally shared globally across species, with species from geographically distant locations hosting similar suites of viruses and viral families.

## Methods

### Sample collection

Sample collection kits and standardized protocols were provided to collaborators to obtain samples of foraging bees from geographically distant locations. For each geographic region, collaborators were instructed to collect western honey bees (*Apis mellifera*) and other common bee species (including both *Apis* and non-Apis bee species) as individuals foraged on flowering plants. Individuals from different bee species were collected in separate tubes to ensure that the species were not mixed. At least 20 individuals were collected onto ice-cold 95% ethanol for each species from each region. Details for each site were recorded, including the GPS coordinates. A summary of different locations and sampled species can be found in Table 1. The samples were then shipped to Pennsylvania State University for processing. We also include samples of *A. mellifera* with deformed wings collected from a colony at the Penn State University apiaries as a control for DWV detection (labeled Positive Control in Table 1).

### Virus purification and nucleic acid extraction

10 individuals from each sample group (representing a specific species collected at a specific location) were placed in individual 2.0 ml microcentrifuge tubes with 150 *μ*l nuclease free H2O and 3–5 sterile glass beads and homogenized in a FastPrep FP120 cell disruptor (Thermo-Fisher Scientific, Waltham, MA, USA) for 45s. 100 *μ*l of the homogenate from each of the 10 individuals was pooled in a common 1.5 ml microcentrifuge tube. The pooled homogenate was then dialyzed using a 0.2 μm cell filter (Corning, Tewksbury, MA, USA) to purify the virus extracts as in Hunter et al. [35] and Liu et al. [36]. A 125 *μ*l aliquot of each sample was treated with a nuclease cocktail (including 14U Turbo DNase I, 25U Benzonase, 20U RNaseI, and 10X DNase buffer) and incubated at 37°C for 1.5 hours to remove all nucleic acid that was not protected by a viral capsid as in He et al. [37]. Nucleic acids (both RNA and DNA) from the purified encapsulated viruses were extracted using a MagMAX Viral Isolation Kit (Thermo-Fisher Scientific, Waltham, MA, USA), according to the manufacturer’s protocol.

### Sequence independent nucleic acid amplification and sequencing

A previously established protocol was used to obtain unbiased random amplification of the extracted nucleic acids [38]. Briefly, the first strand cDNA was synthesized from 30 *μ*l of viral nucleic acid using a random primer design (consisting of a 20 base oligonucleotide sequence followed by a randomized octamer sequence: GACCATCTAGCGACCTCCACNNNNNNNN) in a reverse transcriptase (RT) reaction using a High-Capacity cDNA Reverse Transcription Kit (Thermo-Fisher Scientific, Waltham, MA, USA). To synthesize the second strand, the 19 *μ*l of the initial cDNA was denatured at 95 °C for two minutes and cooled to 4 °C prior to the addition of Klenow fragment DNA polymerase (New England Biolabs, Ipswich, MA, USA) for fragment extension at 37 °C for 60 minutes. Finally, PCR amplification was performed for 40 cycles using the above primer without the randomized octamer sequence (GACCATCTAGCGACCTCCAC) and 2 μl of dsDNA template. The PCR reactions were then purified using a MSB Spin PCRapace Purification Kit (Invitek, Berlin, Germany) using the manufacturer’s standard protocol. Quality and concentration of the samples was assessed using a 2100 Bioanalyzer (Agilent, Santa Clara, CA). The samples were then submitted to the Genome Core Facility at Penn State University for sequencing on the Illumina MiSeq, resulting in 1,056,165 (+/− 105,794) reads at 150 bases per sample (see Supplementary Table 1 for more information on the number of reads and assembled contigs in each sample). Raw data from this Whole Genome Shotgun project have been deposited at DDBJ/ENA/GenBank under the accession PEHZ00000000.

### Data processing

The quality of raw sequencing reads was first manually inspected using FastQC [40]. Illumina adaptor sequences and reads with low quality scores (25) were removed using Trimmomatic v0.30 [41]. To identify novel viruses, the trimmed and filtered sequencing reads were assembled *de novo* using three different programs: Oases v0.2.8/velvet v1.2.10 (N50: 310 nt) [42], Trinity v2.1.0 (N50: 326 nt) [43], and SPAdes v3.7.1 (N50: 315 nt) [44]. The N50 (weighted median length of the contigs) for all assemblies were low (the total read length was 150 nt), so the three assemblies for each sample were merged using the software package Minimus2 (AMOS v3.1.0) [45], resulting in an improved N50 of 699 nt.

The assembled contigs were then subjected to a series of processing steps. To remove contamination and retain a set of candidate contigs likely to be viral in origin, each assembly was screened for homologous sequences from three potential sources of contamination: the human genome (version hg38), all bacterial genomes in the NCBI database, and the honey bee genome (version amel4.5). The total set of contigs was screened for sequence homology to the three possible sources of contamination using BLAST [46]. Contigs with significant hits to any of the contamination sources were then removed from further analyses (see Supplementary Table 1 for the number of contigs matching each source of contamination). To identify the closest known viral sequence homology, the remaining assembled contigs were compared to the all sequences in the NCBI nr database [47] using DIAMOND v0.8.36[48] with an E-value cutoff of 10^-5^. Viral families were assigned using the 2017 report of the International Committee on Taxonomy of Viruses (ICTV) for the classification and nomenclature of viruses [49] (see Supplementary Table 2).

To determine relative abundances of each virus, the trimmed sequence reads from each sample were then mapped to the set of candidate viral contigs identified in that sample using Tophat2 v2.0.9 [50] (see Supplementary Table 2). Contigs with >0.1% of the total reads successfully mapping in that sample were chosen for further analyses (see Supplementary Table 3). For the contigs with >0.1% of the total reads, ORFfinder [51] was used to identify potential open reading frames within the contigs that had significant BLAST hits (E-value <1 × 10^−5^) to known viruses to further characterize the viral contigs from each sample. Viral contigs that showed evidence of a putative RNA dependent RNA polymerase (RdRp) or other replicase proteins were selected for further phylogenetic analyses. Note that while we used a sequence independent amplification approach to generate the libraries for sequencing, amplification bias may still have skewed the total numbers of reads for the viral sequences.

### Phylogenetic analysis

Reference sequences corresponding to the RdRp or other replicase proteins were downloaded from NCBI GenBank [52] for viruses that were closely related to the significant BLAST hit to identify the relationship of the novel viruses discovered here to other known viruses (see Figure 2 for accession number for each sequence). Multiple amino acid sequence alignments were performed using MUSCLE [53] in MEGA7 [54] with default settings. Phylogenetic trees for each novel virus identified in Figure 2 were created using the maximum-likelihood method with 500 bootstrap replications in MEGA7 [54], by applying Neighbor-Join and BioNJ algorithms to a matrix of pairwise distances estimated using a JTT model, and then selecting the topology with the superior log likelihood value.

## Acknowledgements

We would like to thank Mario Padilla, Seymon Mukaku, Frederick Onyengo, Drew Wham and Briana Ezray for assistance with sample collection. This project was funded by a grant from USDA-APHIS (1515-8130-0501-CA) awarded to CMG and a National Geographic Young Explorer’s Grant to ZLF and JTK.

